# Locomotion induces stimulus-specific response enhancement in adult visual cortex

**DOI:** 10.1101/109660

**Authors:** Megumi Kaneko, Yu Fu, Michael P. Stryker

**Affiliations:** Center for Integrative Neuroscience and Department of Physiology, University of California, San Francisco, CA 94143, USA; Singapore Bioimaging Consortium, Agency for Science Technology and Research (A*STAR), Singapore

## Abstract

The responses of neurons in the visual cortex (V1) of adult mammals have long been thought to be stable over long periods. Here, we investigated whether repeated exposure to specific stimuli would enhance V1 visual responses in mice using intrinsic signal imaging through the intact skull and two-photon imaging of calcium signals in single neurons. Mice ran on Styrofoam balls floating on air while viewing one of three different, high-contrast visual stimuli. V1 responses to the stimuli that were viewed by the animal were specifically enhanced, while responses to other stimuli were unaffected. Similar exposure in stationary mice, or in mice in which NMDA receptors were partially blocked, did not significantly enhance responses. These findings indicate that stimulus-specific plasticity in the adult visual cortex depends on concurrent locomotion, presumably as a result of the high-gain state of visual cortex induced by locomotion.

**Significance Statement:** We report a rapid and persistent increase in visual cortical responses to visual stimuli presented during locomotion in intact mice. We first used a method that is completely non-invasive, intrinsic signal imaging through the intact skull. We then measured the same effects on single neurons using 2-photon calcium imaging and found that the increase in response to a particular stimulus produced by locomotion depends on how well the neuron is initially driven by the stimulus. To our knowledge, this is the first time such enhancement has been described in single neurons or using non-invasive measurements.

## INTRODUCTION

It is well documented that visual experience during a critical period in early postnatal life can profoundly alter the responses of neurons in the developing mammalian visual cortex. In contrast, primary sensory cortex in adult mammals has been thought to be “hard-wired” by comparison. While many recent results have demonstrated some degree of activity-dependent plasticity in adult V1 (reviewed by Karmarkar and Dan, 2006; Espinosa and Stryker, 2012; Gilbert and Li, 2012; Medini, 2014; van Versendaal and Levelt, 2016), most reports of adult plasticity find it to be smaller, slower, and qualitatively different from that in early life. Therefore, steps to expand the capacity of the adult brain for plasticity are of great interest because of the relevance to recovery from CNS disease and injury and for maintaining cognitive ability well into the last days of life. Previous studies have increased adult plasticity by various invasive and non-invasive manipulations such as environmental enrichment (Greifzu et al., 2014)), anti-depressant treatment (Maya Vetencourt et al., 2008), inhibitory neuron transplant (Southwell et al., 2010), and others (reviewed by Espinosa and Stryker, 2012). We have shown that exposure to high-contrast visual stimuli during locomotion improved recovery of visual cortical responsiveness from amblyopia (Kaneko and Stryker 2014) and enhanced the plasticity induced in adult cortex by monocular deprivation (Fu et al., 2015). Previous studies using evoked potential recordings through electrodes implanted in the visual cortex reported rapid and persistent increases in responses to repeated stimuli (Frenkel et al., 2006; Cooke and Bear, 2010). We wondered whether the behavioral intervention that we have used in our previous studies, i.e. running + visual exposure (VE), would enhance responses in the intact visual cortex of normal adults.

Here, we used intrinsic signal imaging through the intact skull, a completely non-invasive technique, to investigate whether repeated exposure to specific stimuli would enhance visual responses in adult mouse primary visual cortex (V1). We found that V1 responses to the stimuli that were viewed by the animal during daily running on a freely-moving spherical treadmill were specifically enhanced, leaving responses to other stimuli unaffected. The enhancement was prevented by an NMDA receptor antagonist and persisted for at least a week following cessation of 10 days of stimulus exposure. Similar exposure in mice that were not walking or running did not significantly enhance responses. Longitudinal two-photon Ca^2+^ imaging revealed that the average response magnitude to the exposed orientation was significantly increased, while that to the orthogonal orientation was unchanged. These changes in responsiveness were observed in cells whose initial preferred orientation were close to the experienced orientation and in cells with lower orientation selectivity before exposure, and resulted in an attractive shift of cells’ preferred orientation toward the exposed one and a sharpening of orientation tuning.

### MATERIALS AND METHODS

#### Animals

C57BL/6J (RRID:IMSR_JAX:000664) wild type mice were purchased from Jackson Laboratory (Bar Harbor, ME) and bred as needed, and animals of either sex were used. Animals were maintained in the animal facility at University of California San Francisco and used in accordance with Protocol AN143347 approved by the UCSF Institutional Animal Care and Use Committee. A custom stainless steel plate for head fixation was attached to the skull with dental acrylic under isoflurane anesthesia. The exposed surface of the skull was covered with a thin coat of nitrocellulose (New-Skin, Medtech Products Inc., NY) to prevent desiccation, reactive cell growth, and destruction of the bone structure. Animals were given a subcutaneous injection of carprofen (5 mg/kg) as a post-operative analgesic. All mice were housed in groups of 4–5 and kept under standard conditions (12-h light/12-h dark cycle, free access to food and water) between recordings and daily running on the treadmill.

#### Intrinsic signal optical imaging

Repeated optical imaging of intrinsic signals was performed as described (Kaneko et al., 2008). Five to 7 days after the headplate implantation, the first imaging of intrinsic signals was performed to measure baseline responses. The mouse was anesthetized with isoflurane (3% for induction and 0.7% during recording) supplemented with intramuscular injection of chlorprothixene chloride (2 μg/g body weight) and images were recorded transcranially through the window of the implanted headplate. Intrinsic signal images were obtained with a Dalsa 1M30 CCD camera (Dalsa, Waterloo, Canada) with a 135 mm × 50 mm tandem lens (Nikon Inc., Melville, NY) and red interference filter (610 ± 10 nm). Frames were acquired at a rate of 30 fps, temporally binned by 4 frames, and stored as 512 × 512 pixel images after binning the 1024 × 1024 camera pixels by 2 × 2 pixels spatially. Responses in each mouse were measured with three kinds of visual stimuli (1) horizontal bars drifting upward or downward, (2) vertical bars drifting leftward or rightward, and (3) the contrast-modulated noise movie. They were generated in Matlab using Psychophysics Toolbox extensions (Brainard, 1997; Pelli, 1997), and displayed on a LCD monitor (30 × 40 cm, 600 × 800 pixels, 60-Hz refresh rate) placed 25 cm from the mouse, spanning ~60° (height) × ~77° (width) of visual space. The drifting bar was the full length of the monitor and 2° wide, and it moved continuously and periodically (Kalatsky and Stryker 2003). The contrast-modulated Gaussian noise movie consisted of the Fourier-inversion of a randomly generated spatio-temporal spectrum with low-pass spatial and temporal cutoffs applied at 0.05 cpd and 4 Hz, respectively (Niell and Stryker, 2008). To provide contrast modulation, the movie was multiplied by a sinusoid with a 10-s period. Movies were generated at 60 × 60 pixels and then smoothly interpolated by the video card to 480 × 480 to appear ~60° (height) × ~60° (width) on the monitor and played at 30 frames per second. Each recording took 240 s and was repeated for at least 6 measurements per animal. During the daily session of running + visual exposure (VE), animals were exposed to only one of these 3 stimuli.

#### Analysis of intrinsic signal images

The ROI within V1 was selected on the response magnitude map evoked by visual stimulation. First, the map was smoothed to reduce pixel shot noise by low-pass filtering using a uniform kernel of 5 × 5 pixels. The background area was selected from the area covering ~150 × 150 pixels outside of V1. The ROI was selected by thresholding at 40% above the average background amplitude and the response amplitude was then calculated as the average amplitude of pixels within the ROI.

#### NMDA receptor blockade

The NMDA receptor antagonist 3-(2-carboxypiperazin-4-yl)pro-pyl-1-phosphonic acid (CPP; Tocris Bioscience) (10 mg/kg) or vehicle solution was injected intraperitoneally 1 h before running + VE sessions every day. This dosage was shown to inhibit visual cortical plasticity but not to affect general behavior in adult mice (Sato and Stryker, 2008).

#### Two-photon imaging of Ca^2+^ signals

AAV1-Syn-GCaMP6s (UPenn Vector Core) was injected stereotaxically into the primary visual cortex (V1) around P30. Approximately 2 weeks after virus injection, a custom titanium head plate with a 3 mm-diameter window was fixed to the skull and a round glass coverslip (3mm diameter) was cemented over a craniotomy made over V1, under isoflurane anesthesia (3% induction, 1.5–2% maintenance) supplemented with subcutaneous analgesics injections. Five to 7 days after window implantation, baseline Ca^2+^ responses were recorded under the same anesthesia as for the intrinsic signal imaging. Two to 3 days after recording baseline responses, each animal underwent daily sessions of running + VE.

Drifting sinusoidal grating stimuli were generated using the Psychophysics Toolbox extensions in MatLab (MathWorks) and displayed on a LCD monitor (Dell, 30×40 cm, 60Hz refresh rate) placed 25 cm from the mouse. Each trial stimulus consisted of a 3 s grating (0.05 cycle per degree, 1 Hz temporal frequency) followed by a 3 or 4 s of blank period of uniform 50% grey. Eight drifting directions in 45° steps presented at random sequence were repeated 6 times per recording set.

Imaging was performed using a Sutter Movable Objective Microscope and a Chameleon ultrafast laser (running at 940 nm), controlled by ScanImage (http://scanimage.org). Images were collected at 5 Hz, 512 × 256 pixels covering ~250 μm × 250 μm at the depth of 150–300 μm from the dura surface.

#### Ca^2+^ signal analysis

Regions of interest (ROIs) corresponding to visually identifiable cell bodies were selected manually, and the nuclear region, which is devoid of fluorescence, was excluded. Cells with overlapping ROIs were excluded from the analysis. The fluorescence time course of each cell was obtained in ImageJ by averaging all pixels within the ROI. Further analyses were performed by a custom written Matlab program (Fu et al., 2014). Briefly, ΔF/F_0_ was calculated as (F-F_0_)/F_0_, where F_0_ is the baseline fluorescence signal averaged over a 1-s period immediately before the start of visual stimulation. Visual responses were measured for each trial as ΔF/ F_0_, averaged over the last 2 s of the stimulus period. Neurons were considered visually responsive when fluorescence changes were significantly related to stimulus (ANOVA across blank and 8 direction stimulus periods, P < 0.01) (Ohki et al., 2005), with an average ΔF/F_0_ at preferred orientations greater than 25%. The preferred orientation (*O*_*prf*) was determined by fitting the tuning curve with the sum of two Gaussians. The orientation selectivity index (OSI) was computed for responsive cells as (R_pref_ − R_ortho_)/ (R_pref_ + R_ortho_), where R_pref_ and R_ortho_ are the response amplitudes at the preferred and the orthogonal orientation, respectively (Niell and Stryker, 2008).

#### Statistical analyses

Data were presented as mean ± SEM, mean ± s.d., or cumulative frequency distributions unless otherwise indicated. Statistical methods employed were stated in the result section and/or figure legends. Statistical analyses were performed using Prism 6 (GraphPad Software, CA) or Matlab (MathWorks, MA).

### RESULTS

#### Response enhancement measured by intrinsic signal imaging

Each animal was acclimated to the running setting by the experimenter’s handling and being placed on the Styrofoam ball floating on air for approximately 15 min a day for 4–5 days. Baseline intrinsic signal responses were then measured to three different, high-contrast visual stimuli: drifting horizontal bar, drifting vertical bar, and contrast-modulated noise movie (Figure 1A). Starting 2–3 days later, animals were allowed to run on Styrofoam balls while viewing one of those three visual stimuli 60 min per day for 10 days, as described previously (Kaneko and Stryker, 2014). To track the change in visual cortical responses to these visual patterns, we repeated intrinsic signal recordings on day 5 and day 10, and then on day 17, one week after stopping the daily running + visual exposure (VE) sessions (Figure 1A). The parameters for the drifting bars were chosen to elicit strong responses in V1 based on our previous report (Kalatsky and Stryker, 2003). The visual stimuli also included the contrast-modulated stochastic noise matched to the spatiotemporal frequency response of the mouse V1 neurons because it drives nearly all cells in the primary visual cortex to some extent (Niell and Stryker, 2008).

**Figure 1.**
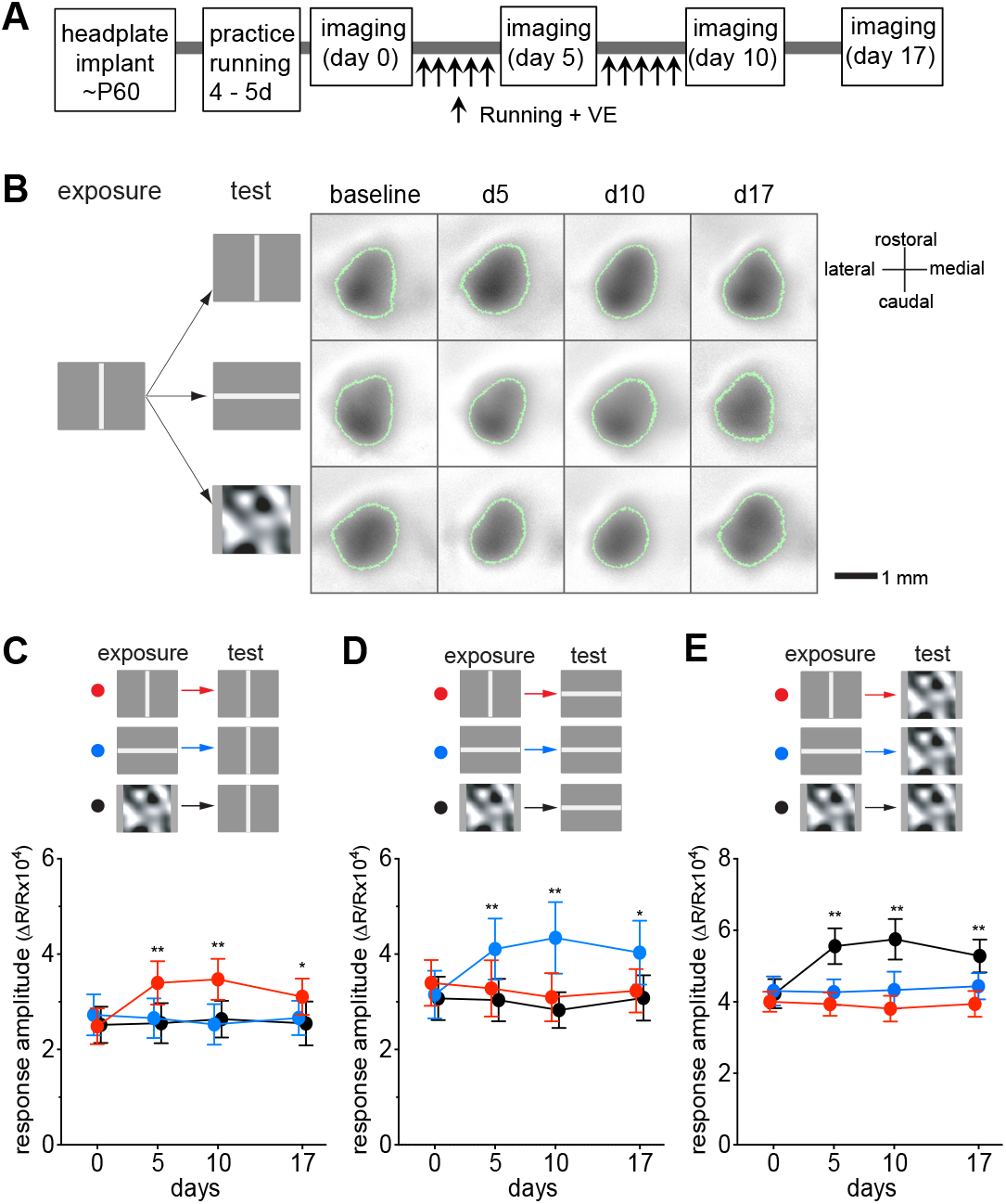
Preferential potentiation of visual cortical responses to the visual stimuli experienced during locomotion. **A.** Experimental schedule. During daily running from day 1 through day 10, mice were presented with either drifting vertical bars, drifting horizontal bars, or contrast-modulated noise, indicated as “exposure” in B-E. In every animal, visual cortical responses to all three visual stimuli were examined, indicated as “test” in B-E. **B.** Examples of intrinsic signals of a mouse that viewed drifting vertical bars during the daily running. Green lines encircling the response area indicate the ROIs from which response averages were calculated. **C-E**. Time course of change in intrinsic signal responses to drifting vertical bars (C), drifting horizontal bars (D), or contrast-modulated noise (E), in animals exposed to the vertical bars (red circles, n = 6), the horizontal bars (blue circle, n = 6), or the noise (black circles, n = 6). Each point in graphs represents mean ± SEM of 6 animals; at each time point, at least 5 measurments of the averaged response over the ROI were averaged for each animal. **P < 0.01, *P < 0.05 compared with the baseline (day 0) response; two-way ANOVA followed by multiple comparisons with Bonferroni’s correction.

Figure 1B shows examples of intrinsic signal images recorded over time from a mouse that viewed drifting vertical bars during daily running on the ball, revealing a modest increase in the response to the vertical bars and little change in responses to horizontal bars or the noise movie. As a group, mice that viewed vertical bars during running showed a modest but significant increase in response to the same stimulus (red circles in Figure 1C) (baseline: 2.49 ± 0.84, d5: 3.40 ± 1.02, d10: 3.47 ± 0.96; two-way ANOVA _(6,36)_ = 6.6, P < 0.001, for the interaction effects of visual stimuli and days; F_(3,36)_ = 2.1, P = 0.11 for the effect of days; P < 0.01 on d5 and d10 compared to baseline) but responses to other stimuli were unchanged (red circles in Figure 1D, response to horizontal bars baseline: 3.39 ± 1.08, d5: 3.28 ± 1.32, d10: 3.09 ± 1.13; and in Figure 1E, response to the noise movie baseline: 4.00 ± 0.63, d5: 3.93 ± 0.73, d10: 3.81 ± 0.81; P > 0.05 on d5 and d10 vs. baseline). Likewise, mice that viewed horizontal bars during running demonstrated a modest but statistically significant increase in responses when tested with the same visual stimulus but not to stimuli that they had not experienced (blue circles in Figure 1C–E; responses to horizontal bars, baseline: 3.15 ± 1.12, d5: 4.11 ± 1.43, d10: 4.34 ± 1.68; vertical bars; baseline: 2.73 ± 0.96, d5: 2.66 ± 0.93, d10: 2.53 ± 0.95; noise movie: baseline: 4.30 ± 0.91, d5: 4.27 ± 0.81, d10: 4.33 ± 1.16; two-way ANOVA F_(6,36)_ = 6.83, P < 0.001 for the interactions of visual stimuli and days, F_(3,36)_ = 4.84, P = 0.006 for the effects of days). A similar change in responsiveness to the exposed stimulus was observed in mice that viewed contrast-modulated noise movie during running (black circles in Figures 1C–1E; responses to noise movie, baseline: 4.22 ± 0.91, d5: 5.56 ± 1.12, d10: 5.75 ± 1.27; responses to vertical bars: baseline: 2.52 ± 0.86, d5: 2.55 ± 0.95, d10: 2.64 ± 0.86; responses to horizontal bars, baseline: 3.07 ± 1.02, d5: 3.04 ± 1.00, d10: 2.83 ± 0.85; two-way ANOVA F_(6,36)_ = 12.28, P < 0.001 for the interaction effects of visual stimuli and days, F_(3,36)_ = 10.29, P < 0.001 for the effects of days). These enhancements had peaked by day 5 with only a slight, insignificant increase on day 10, and they persisted for at least 7 days after terminating sessions of running + VE (d17 in Figure 1C–E).

One hr/day of running + VE was sufficient to produce a saturating effect. Increasing the duration of daily running + VE from 1 to 2 to 4 hrs did not produce significantly greater enhancement (Figure 2).

**Figure 2.**
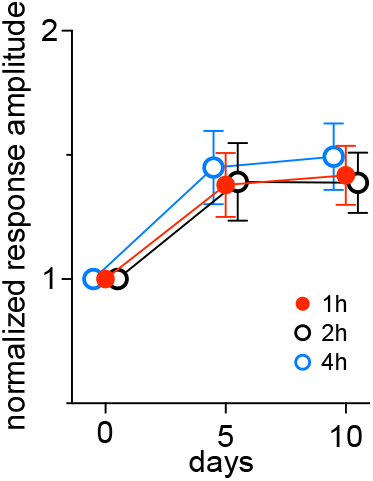
Increasing the duration for running + visual exposure did not further increase response enhancement. Animals were placed on the spherical floating treadmill daily for 2 hours (2h: n = 4) or for 4 hours (4h: n = 4) while viewing a vertically-oriented bar drifting horizontally. Responses in V1 were measured using intrinsic signal imaging and were normalized to baseline value (day 0). Error bars represent mean ± s.d. Data for 1h were from Figure 1C (n = 6). Changes after 5 days and 10 days of running + visual exposure were not significantly different among 3 groups (two-way ANOVA).

These observations are similar in magnitude to those of the report by Frenkel et al. (2006), which found an approximate 40% increase in visually evoked potentials (VEP) through electrodes chronically implanted into V1 of mice older than P60 after short periods of exposure to specific visual stimuli.

#### Response enhancement depends on NMDA receptor activation

It has been shown that NMDA receptor activation is required for experience-dependent enhancement in responses in adult visual cortex (Sato and Stryker, 2008; Sawtell et al, 2003; Frenkel et al, 2006). We examined whether the enhancing effects of running + VE require NMDA-receptor activation. We inhibited partially NMDA receptors by daily systemic administration of the competitive antagonist CPP (Figure 3A). Running behavior was indistinguishable between vehicle- and CPP-treated animals (% of time spent moving, P = 0.877 for treatments, 2 way ANOVA, Figure 3F; average velocity of movement, P = 0.861 for treatments, data not shown). Intrinsic signal responses to the visual stimuli that the animals viewed during daily running were enhanced in the vehicle-treated control mice just as in untreated animals (Figure 3B-D). In contrast, CPP treatment completely blocked these changes over the course of 10 days (Figure 3B, baseline: 2.81 ± 0.96, d5: 3.00 ± 1.03, d10: 3.08 ± 1.05; Figure 3C, baseline 3.40 ± 1.18, d5: 3.48 ± 1.22, d10: 3.60 ± 1.17; Figure 3D, baseline 4.03 ± 0.83, d5: 4.45 ± 1.00, d10: 4.70 ± 1.16; P > 0.05 on d5 and d10 vs. baseline). This CPP treatment regimen did not change cortical responsiveness in mice that viewed a blank screen of 50% grey during running (Figure 3E).

**Figure 3.**
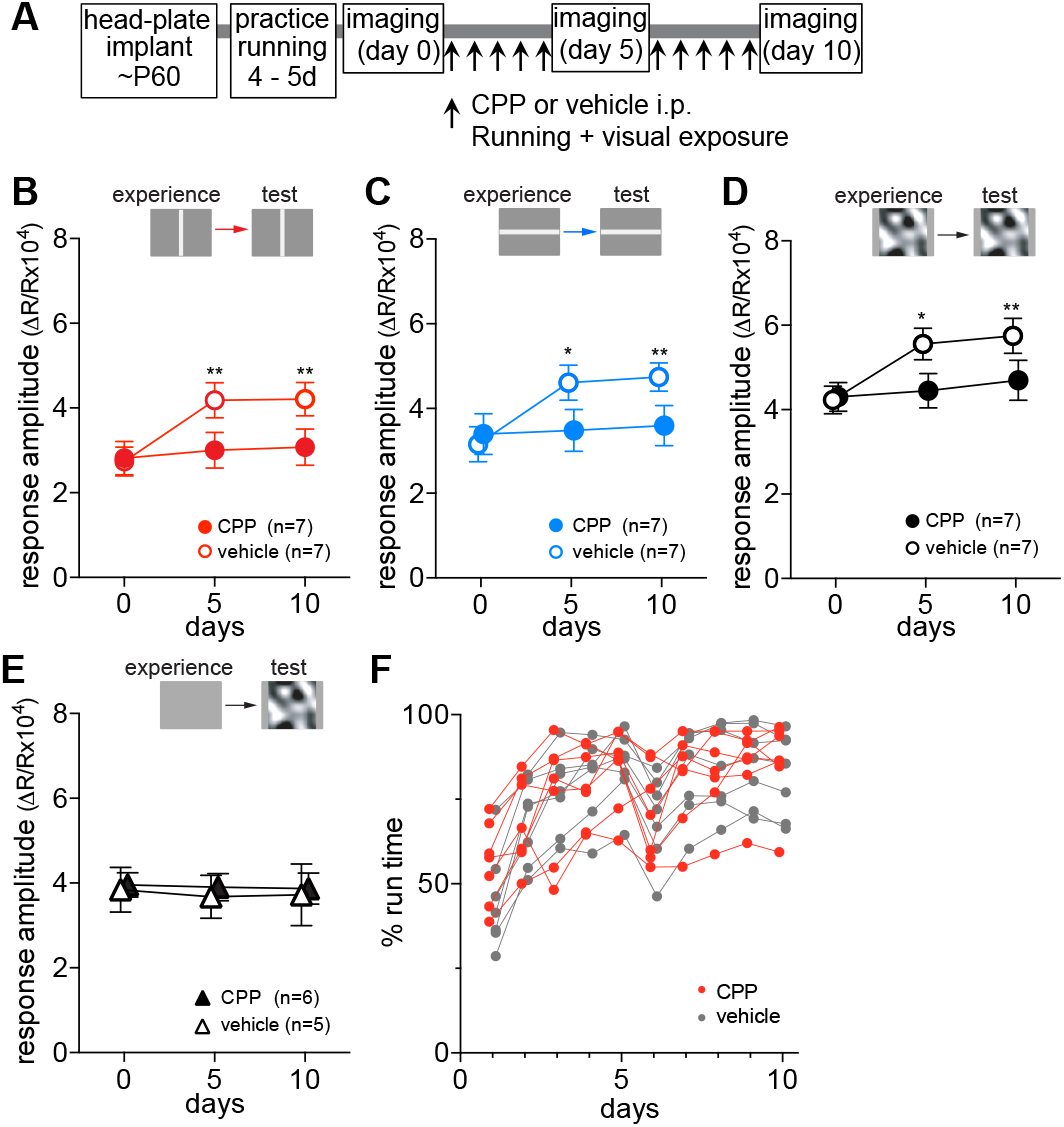
Response potentiation depends on NMDA receptor activation. **A.** Experimental schedule. **B-D.** Changes in responses to drifting vertical bars (B), to drifting horizontal bars (C), or to contrast-modulated noise (D) in mice that experienced same visual patterns during daily running. Closed circles and open circles represent CPP-treated (n = 7) and vehicle-treated (n = 7) animals, respectively. **E.** Lack of effects of partial NMDA receptor blockade on stability in response magnitude of intrinsic signals over the period of 10 days in animals that were exposed to an uniform 50% grey screen during daily running. **F.** Percentage of time that each mouse moved > 1 cm/s on the Styrofoam ball during daily running + VE in CPP-treated (red circles) and vehicle-treated (grey circles) groups shown in B. Data in B-E are presented as mean ± SEM. *P < 0.05, **P < 0.01 compared with baseline (day 0) within the group; two-way ANOVA followed by multiple comparisons with Bonferroni correction.

#### Locomotion is required for response enhancement

The findings above reveal that response enhancement depends on visual stimulation, in that looking at a blank screen during running had no apparent effect (Figure 3E), and is stimulus-specific, affecting only the stimulus viewed during locomotion (Figure 1). Our previous study had demonstrated that both locomotion and visual stimulation were necessary for the recovery of cortical responses to normal levels in amblyopic adult mice in which one eye had been occluded through the critical period for 5 months (Kaneko and Stryker, 2014). Recovery did not take place with visual stimulation in the home cage, where sustained running was not possible, or with locomotion facing a constant grey screen. But what is the role of locomotion for enhancement in normally reared mice?

To determine whether locomotion is necessary for the enhancement we observed above, as it is for recovery from deprivation, we severely restricted the animal’s running by almost completely shutting off the air supply to the system that makes the Styrofoam ball float. To acclimatize animals to restricted locomotion, we started the air flow at ~50% of normal and gradually decreased it to almost zero over several days. Some mice (approximately 40% of animals that got started practice) failed to be still at all after 10 days of training and were excluded from the experiments. The movement of the Styrofoam ball was monitored as described (Niell and Stryker, 2010). The animals were trained to sit still on the movement-restricted ball for 8–10 days, approximately 20 min/day. After recording the baseline intrinsic signal responses on day 0, animals were placed on the movement-restricted ball while viewing drifting vertical bars 60 min/day for 10 days, and intrinsic signal recordings were repeated on day 5 and 10 (Figure 4A). In contrast to the condition in which the animals were allowed free locomotion, cortical responses to the experienced stimulus were not significantly enhanced when the animals’ movement was restricted (Figure 4B, responses to vertical bars, baseline: 2.03 ± 0.58, d5: 2.03 ± 0.43, d10: 2.05 ± 0.48; P > 0.05, repeated measure ANOVA; compare to solid pink circles showing enhancement with locomotion).

**Figure 4.**
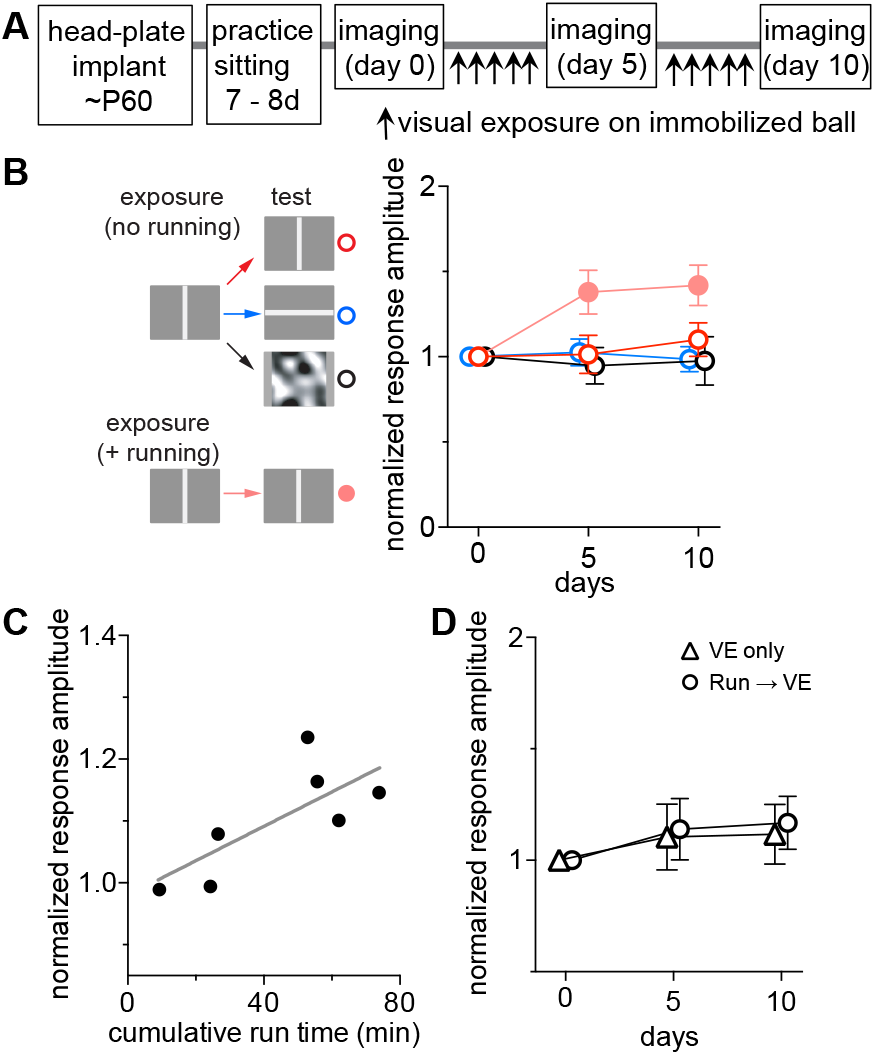
Restricting locomotion reduces stimulus-specific enhancement of response magnitude. **A.** Experimental schedule. **B.** Changes in response magnitudes in animals that viewed drifting vertical bars daily while on the Styrofoam ball the movement of which was restricted. Intrinsic signal imaging was performed in each animal to measure cortical responses to three different visual stimuli, vertical bars (red), horizontal bars (blue), and contrast-modulated noise (black). Change in response amplitude in animals that moved freely on the ball while viewing drifting vertical bars, shown in solid pink circles, data from Figure 1C. Response amplitudes were normalized to baseline (day 0) and shown as mean ± s.d. *P < 0.05, **P < 0.01 between the free run group and restricted move group; two-way ANOVA followed by multiple comparisons with Bonferroni correction. **C.** Change in response amplitude as a function of total duration of movement in animals that were placed on balls with restricted movement. Each point shows data from individual animals in B. The y-axis represents normalized responses to drifting vertical bars (the exposed visual stimulus) at day 10. The x-axis represents 10-day cumulative duration that the trackball was considered moving (> 1 cm/s). R^2^ = 0.58; slope 0.0029 ± 0.0011 (95% CI: 4.39e^−0.0005^, 0.0057); slope deviation from zero P = 0.047; least-squares linear regression. **D.** Visual experience while not engaging in running did not enhance V1 responses. One group of animals were allowed to run freely on a floating Styrofoam ball while viewing a blank grey screen for 1 hour daily, followed immediately by being returned to home cage and exposed to a drifting vertical bar for 1 hour, on day 1 through day 10 (Run → VE; n = 4). Another group of mice were exposed to the same visual stimulus in the home cage without preceding running (VE only; n = 4). Visual exposure was done while the animals were placed in the housing cage made of clear polycarbonate that were surrounded by four monitors, one on each side, displaying the visual stimulus. Responses to the experienced stimulus (vertical bar) were recorded on day 0 (baseline), day 5, and day10. Data are normalized to the baseline value and presented as mean ± s.d. Response magnitude on day 5 or day 10 were not significantly different from baseline in either group (two-way ANOVA).

Although the movement of the ball was restricted in these experiments, it was not completely immobilized, and we noticed that some mice moved it more than others. The 4 animals that succeeded in moving the ball for a total of nearly an hour over the 10 hours of exposure all showed some degree of enhancement (Figure 4C). Indeed, the changes in response magnitude to the experienced stimulus were positively correlated with the duration of movement (Figure 4C; slope = 0.0029 ± 0.0011, P = 0.047 for deviation from zero; R^2^ = 0.58; least squares linear regression). Locomotion speed in the movement-restricted condition, while classified as “moving”, was much lower (1-3 cm/sec) compared to that in freely-moving condition.

It has been shown that a beneficial effect of acute aerobic exercise in young human volunteers (improvement on an orientation discrimination task after 30 min on a stationary bicycle) can last at least 30 min after stopping the exercise (Perini et al., 2016). To test whether locomotor activity confers such benefit in mice, we allowed animals to run for 1 hr while viewing a grey screen and then returned them to the home cage and exposed them to a high contrast visual stimulus immediately thereafter. A second group had only the exposure in the home cage without the prior running. Intrinsic signal responses did not change significantly in either group, confirming that the visual stimulation must be concurrent with locomotion for enhancement (Figure 4D).

Taken together, these observations indicate that stimulus-specific enhancement is dependent on a cortical state that is induced by locomotion and not merely on repeated exposure to the stimulus.

#### Response enhancement measured in single cells

To begin to understand cellular mechanisms of the stimulus-specific response enhancement observed in intrinsic signal imaging experiments, we examined changes in response properties of individual neurons in layers 2/3 of the primary visual cortex using two-photon imaging of Ca^2+^ signals. Two weeks after injecting AAV1-hsyn-GCaMP6s stereotaxically into the primary visual cortex on ~P45, we implanted the chamber for head fixation containing the imaging window centered the injection site, and then gave the mice 3-4 days of practice standing and running on the Styrofoam ball for 15 min/day. We then made baseline recordings of the calcium responses of many single cells to gratings drifting in 8 different directions while the mice were anesthetized. After an additional 2-3 days, the mice were placed for 60 min daily on the floating Styrofoam ball while viewing one of three stimuli: gratings oriented at 45 degrees drifting back and forth (G45, 2 mice), drifting gratings oriented at 135 degrees (G135, 2 mice), or a blank (50% grey) screen (Control, 2 mice). After 5 days of running + VE, the calcium recordings were repeated so as to measure responses in the same cells that were studied at baseline (Figure 5A).

**Figure 5.**
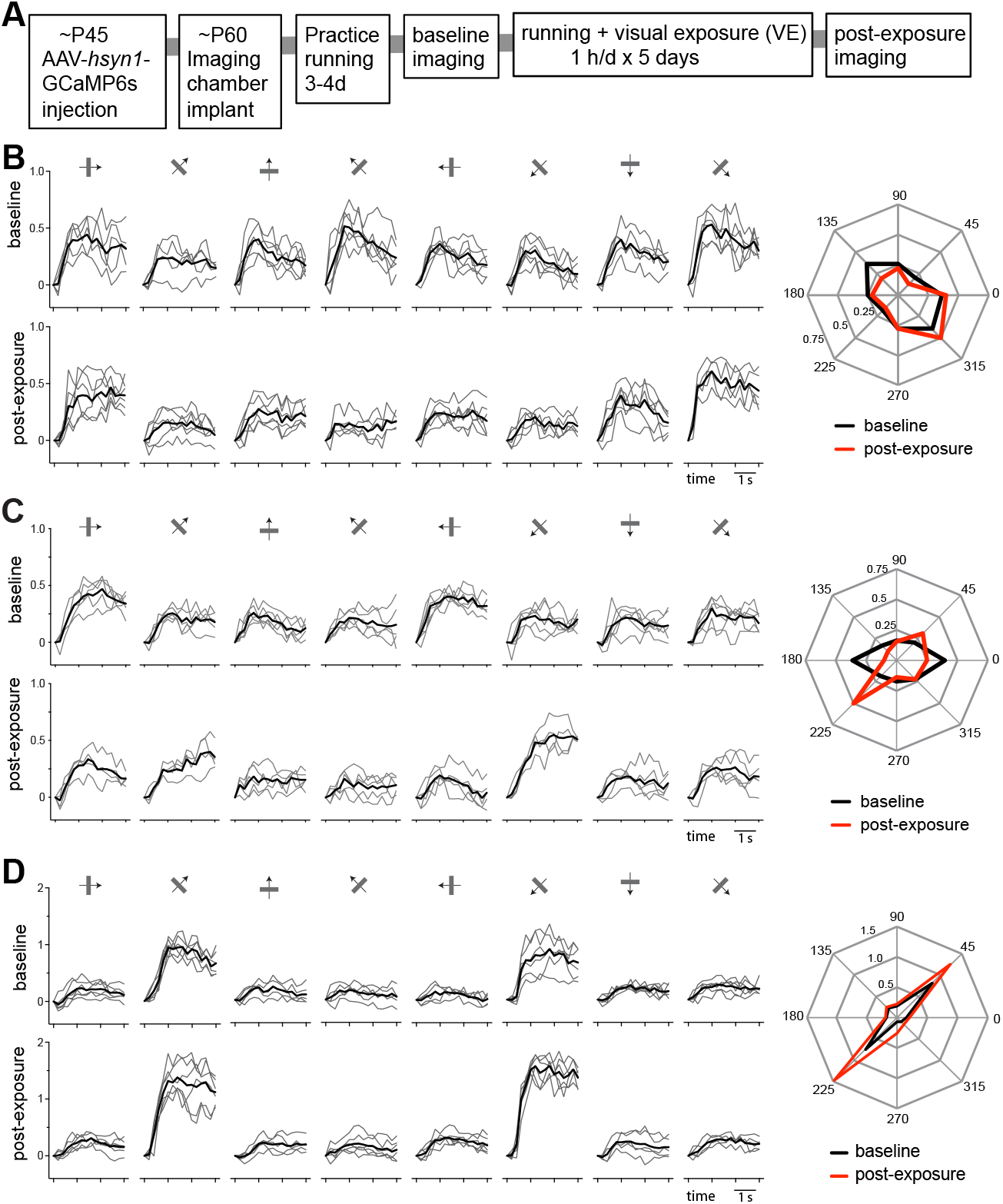
Tracking changes of calcium responses in individual cells. **A.** Experimental schedule. **B–D.** Examples of calcium signal traces in response to drifting gratings of eight different directions in a cell in a control animal (B) and two cells in a experimental animal (C and D, both are in G45 group). Left panels: Upper: baseline recording; Lower: post-exposure recording. Thin grey lines: individual trial; Black lines: mean of 5-6 individual trials. Right panels: Polar plots of averaged responses in baseline (black) and post-exposure (red) recordings. Responses to each direction were averaged over the last 2 seconds of stimulus presentation.

Of the total of 838 cells that were identifiable in both baseline and post-exposure recordings (control: 324, G45: 286, G135: 228 cells), we selected cells with peak ΔF/F ≥ 0.25 in both recordings as responsive cells for further analyses (control: 192, G45: 172, G135: 166 cells). Figure 5B–D show examples of responses at baseline and post-exposure in the control and G45 groups. The cell shown in Figure 5B from a control animal had moderately tuned baseline responses that did not change in either amplitude or selectivity between the two recording sessions (preferred orientation *(O_prf)*: 140.1 vs. 144.9; OSI: 0.40 vs. 0.48; Gaussian fit peak: 0.44 vs. 0.49; baseline vs. post-exposure). The cell shown in Figure 5C from a G45 animal had moderately tuned baseline responses (OSI: 0.38) similar to that of a control cell shown in 5B. Its post-exposure responses to the exposed orientation *(O_exp)* was increased (0.197 vs. 0.406) with little change in responses to the orthogonal orientation (0.198 vs. 0.163) or overall responsiveness (0.236 vs. 0.225; AF/F averaged for all orientations), resulting in a shift of its *O*_*prf* toward the *O*_*exp* (1.4 vs. 34.4). Another example cell from the same G45 mouse, shown in Figure 5D, was initially highly selective for the *O*_*exp* at baseline (*O_prf:* 46.6, OSI: 0.76), and the responses after exposure to that orientation were increased (0.78 vs. 1.35) with little change in responses to the orthogonal orientation (0.16 vs. 0.20), resulting in further sharpening of orientation tuning (*O_prf:* 43.9; OSI 0.82).

On average, cells in running + VE groups significantly increased their responses to the exposed orientations (P < 0.01; one-way ANOVA followed by multiple comparisons with Bonferroni corrections) while their responses to the orthogonal orientations were unchanged (P > 0.05, vs. control) (Figures 6A,B,C). Cells in control animals showed no statistically significant changes in responses to either orientation (Figure 6A,B,C).

**Figure 6.**
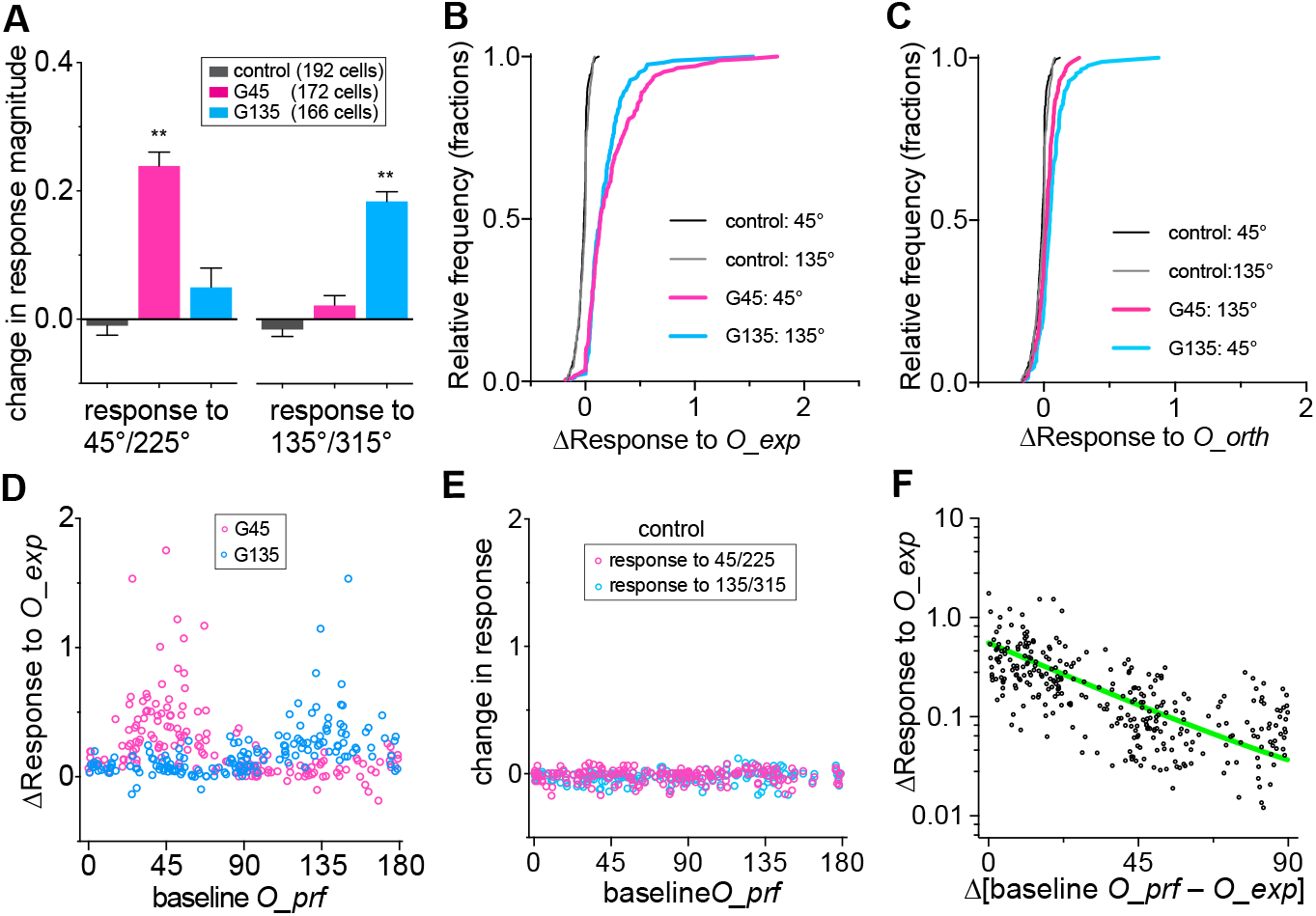
Changes in calcium response magnitudes after running + visual exposure. **A.** Changes in calcium signals in response to gratings of the exposed orientation and the orthogonal orientation. Bars represent mean ± SEM. **P<0.001, *P<0.01 compared with respective control group; one-way ANOVA followed by multiple comparisons with Bonferroni correction. **B, C.** Cumulative frequency distributions of changes in response magnitude to the exposed orientation during running (*O_exp,* E) and the orientation orthogonal to *O_exp* (*O_orth:* F). **D.** Change in responses to orientations that were exposed during locomotion (*O_exp*), plotted against each cell’s baseline preferred orientation (*O_prf*). Each circle represents simple difference in response magnitudes between post-exposure and baseline in a single cell. **E.** Differences in response to gratings at 45°/225° and 135°/315° between the first (baseline) and the second (post-session) recordings plotted against each cell’s baseline *O_prf* in the control group. Each circle represents simple difference in response magnitudes between post-session and baseline in a single cell. **F.** Changes in responses to *O_exp,* shown in (D), were re-plotted to the distance of each cell’s baseline O_prf from *O_exp.* Data from two experimental groups were pooled. Green line represents one phase decay fit; R^2^ = 0.399.

In spite of the prominent increase in average responses to the exposed orientations, changes in individual cells in the experimental groups varied widely. For both G45 and G135 experimental groups, the change in response to *O_exp* depended strongly on each cells’ *O_prf* at baseline (Figure 6D); whereas cells in control mice showed negligible changes in response magnitudes to the two experimental orientations between recording sessions (Figure 6E). The cells initially selective for orientations closer to *O_exps* in the experimental mice showed much larger increases in responses to that orientation than did other cells (Figure 6F, R^2^ = 0.35, least-squares linear regression; R^2^ = 0.40 for non-linear fit exponential one-phase decay).

These findings suggest that the response to a particular stimulus is enhanced by locomotion to an extent that depends on how well the neuron responds to that stimulus at baseline. To test this notion, we plotted the degree of enhancement as a function of the baseline responses (Figure 7). In both experimental groups there was a highly significant relationship between baseline response to the exposed stimulus and enhancement. Responses to the stimuli that were not exposed were not enhanced.

**Figure 7.**
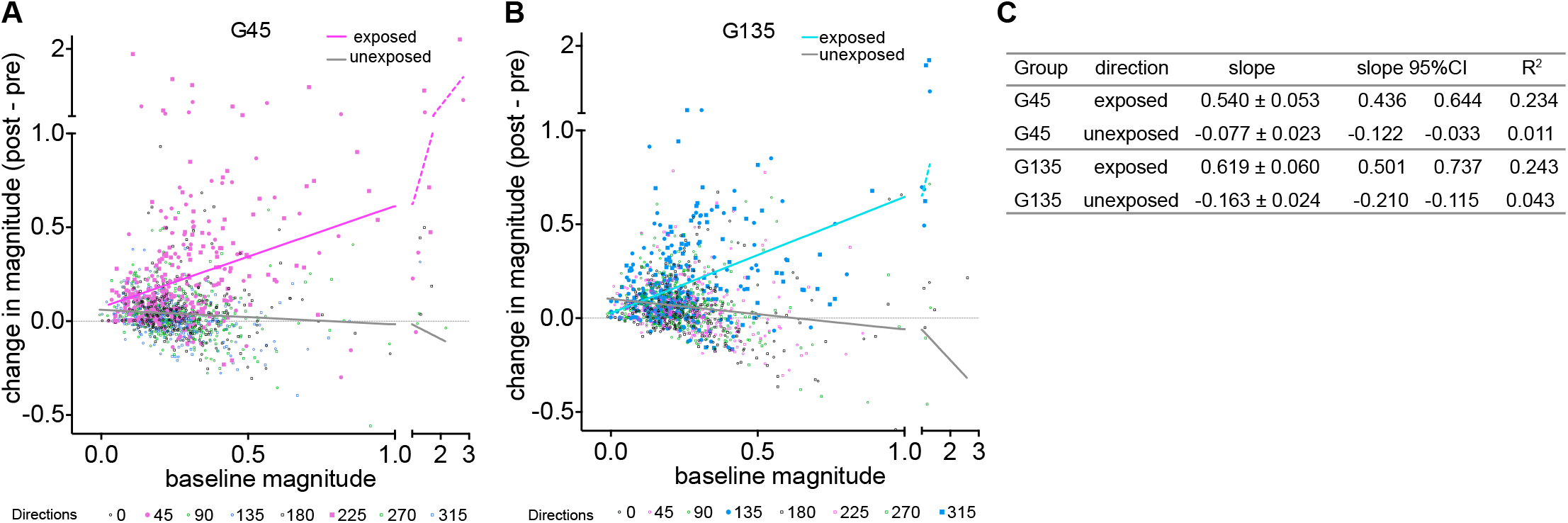
Relationship between baseline Ca^2+^ responses and change in response magnitudes at each direction of drifting grating stimuli. **A, B.** Data from each experimental group were plotted separately (A: G45 group, B: G135 group). Lines represent least squares linear regression, analyzed for the exposed 2 directions and other 6 directions separately. **C.** Statistical data for linear regression. All slopes showed statistically significant deviation from zero (P < 0.0001).

As a result of increase in the response magnitude to the exposed orientation with little change in responses to other orientations, the preferred orientation of cells whose baseline preferred orientations were close to the one exposed shifted toward the exposed orientation (Figure 8A,B,D,E note the flattening of the pre-versus post-exposure curves near *O_exp* in the G45 and G135 groups). Cells in control animals showed no significant changes in their preferred orientations (Figure 8C,D,E). Figure 8F,G compare the shifts in *O_prf* between control and experimental groups.

**Figure 8.**
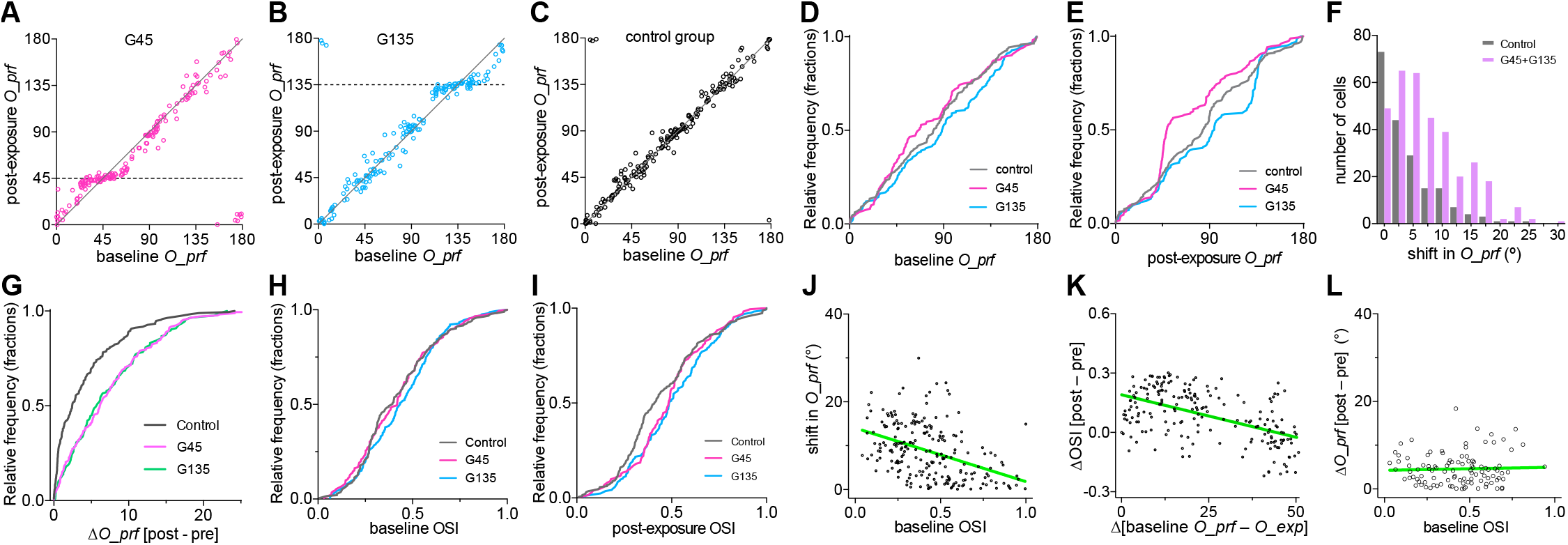
Changes in preferred orientation and orientation selectivity after running + visual exposure. **A-C.** Preferred orientation (*O_prf*) in individual cells at baseline (x-axis) and post-exposure (y-axis) in the G45 (A), G135 (B), and control (C) groups. **D.** Cumulative frequency distribution of preferred orientations (*O_prf*) at baseline. **E.** Cumulative frequency distribution of *O_prf* after exposure to specific orientations of drifting gratings during daily running. **F.** Frequency distributions of shift in *O_prf*, summarized from data shown in A-C. Data from two experimental groups were pooled. P < 0.001 between control and experimental groups, K-S test. **G.** Cumulative frequency distribution of shift in *O_prf* in individual cells. **H.** Cumulative frequency distribution of orientation selectivity (OSI) at baseline. **I.** Cumulative frequency distribution of OSI after the exposure to specific orientations of drifting gratings during daily running. **J.** Inverse correlation between baseline OSI and shift in *O_prf* in experimental groups. Green line represents linear regression, R^2^ = 0.18; slope: −12.21 ± 1.78 (95% CI: −15.71, −8.72), slope deviation from zero P < 0.0001. **K.** Increase in orientation selectivity when the cells’ baseline *O_prf* was closer to *O*_*exp.* The green line represents linear regression; R^2^ = 0.28; slope: −0.0043 ± 0.0005 (95% CI: −0.0053, −0.0033); slope deviation from zero, P < 0.0001. **L.** Shift in *O_prf* had no relationship to baseline OSI in cells in which baseline *O_prf* were more than 50° apart from the exposed orientations. R^2^ = 0.0015, slope: 0.703 ± 1.807 (95% CI: −2.89, 4.29), slope deviation from zero: P = 0.698, least-squares linear regression. For analyses in I and J, we selected cells in which baseline *O_prf* was within 50° from *O_exp* and pooled G45 and G135 (n = 199).

The orientation selectivity index (OSI), calculated as the ratio of response at the preferred to that at the orthogonal orientation, is an additional measure to describe orientation tuning. While distributions of OSI were indistinguishable between groups at baseline (Figure 8H), the OSIs of experimental animals were slightly shifted toward higher values after exposure because of the changes in response amplitudes (Figure 8I, P = 0.002 control vs. G45, P = 0.004 control vs. G135, Kolmogorov-Smirnov test). In cells initially selective for orientations within 50 degrees of *O_exp*, the magnitude of shift in *O_prf* was inversely correlated with the base-line OSI (Figure 8J, R^2^ = 0.18, least-squares linear regression). In this population of cells, increase in OSI was inversely correlated with the difference between baseline *O_prf* and *O_exp* (Figure 8K, R^2^ = 0.28, least-squares linear regression). In cells with baseline *O_prf* further than 50 degrees from *O_exp,* there was no relation between baseline OSI and shifts in preferred orientation (Figure 8L).

These findings reveal that single cells that are driven well by the stimulus presented during locomotion dramatically increase their responses to that stimulus, on average by about 20% and many by 50% or more, with little or no change in other cells (Figure 6A,B). These changes in response magnitude have the effect of slightly shifting the preferred orientations of those cells toward the one exposed, at least as assessed from fits to tuning curves derived from responses to stimuli presented in 45-degree steps. Responses of neurons in control animals exposed to a grey screen during locomotion are, as expected, stable in both magnitude and selectivity (Figures 6C,8C). Results from single cell analysis are thus consistent with those measured using intrinsic signals: response enhancement by locomotion and is stimulus-specific.

### DISCUSSION

Many experiments have found that responses in the primary visual cortex (V1) in adult mammals are stable over long periods. In the present study, using non-invasive repeated imaging of intrinsic signals, we found that the daily exposure of adult mice to high contrast visual stimuli in animals allowed to move freely on a spherical treadmill enhanced the responsiveness of V1 to those stimuli, leaving responses to other stimuli unchanged. This enhancement depended on NMDA receptor activation. Most strikingly, the enhancement to the specific stimuli presented depended on the animals’ locomotion, presumably reflecting the high-gain state into which locomotion places mouse V1 (Niell and Stryker, 2010). Repeated imaging of Ca^2^+ signals in single cells confirmed the stimulus-specificity of the enhancement of V1 responses at the cellular level. They revealed that Ca^2+^ responses to the orientation that was viewed by animals during running was increased while the responses to other orientations were not, a change that produced an attractive shift in preferred orientation toward the one that was viewed.

### A circuit responsible for stimulus-specific enhancement during locomotion

The present finding that the cortical state produced by locomotion is required for enhancement is consistent with our previous observations that recovery of V1 responses from prolonged monocular deprivation was greatly augmented by high-contrast visual stimulation while animals were engaged in locomotion (Kaneko and Stryker, 2014). This recovery depended on a subcorticocortical circuit that increases responsiveness of mouse V1 (Lee et al., 2014; Fu et al., 2014). The circuit originates in ascending projections of the midbrain locomotor region (MLR) to the horizontal limb of the nucleus of the diagonal band of Broca, which sends cholinergic projections to activate vasoactive intestinal peptide containing (VIP) cells in V1 during locomotion. The VIP cells inhibit somatostatin-expressing (SST) GABAergic cells, disinhibiting the excitatory neurons and increasing their responses (Pfeffer et al., 2013; Fu et al., 2014). In general terms, such changes in excitation/inhibition balance are expected to enhance activity-dependent plasticity in adult V1 (Harauzov et al., 2010; reviewed by Bavelier et al., 2010 and Takeshian and Hensch 2013). Experiments inducing loss of function (using tetanus toxin) and gain of function (using optogenetic activation) of VIP cells in mouse V1 revealed that the activity of the VIP-SST disinhibitory circuit was necessary and sufficient in adult V1 to facilitate recovery from amblyopia caused by prolonged MD and to enhance ocular dominance changes by short periods of MD that would otherwise have been ineffective (Fu et al., 2015). It therefore is likely that the same disinhibitory circuit is responsible for the stimulus-specific enhancement in V1 responses that we report in the present study

How might the engagement of this disinhibitory circuit with its transient resetting of E/I balance produce the stimulus-specific enhancement of V1 responses? The present findings from longitudinal Ca^2+^ imaging of single cells may shed light on the cellular mechanisms. On average considered as a single population, V1 neurons showed increased Ca^2+^ responses to the experienced orientation (Figure 6A) while responses to non-experienced orientations were unchanged, consistent with the result at the whole V1 level observed in intrinsic signal imaging. However, the changes at the individual cell level were different from cell to cell. In particular, nearly all of the increase in response to the experienced orientation was in cells in which the baseline preferred orientation was close to the experienced orientation and not in those in which preferred orientation was different (Figure 6B). Although the mouse lacks orientation columns, L2/3 pyramidal neurons in V1 with similar orientation selectivity preferentially form synapses with one another; that is, V1 is organized into subnetworks defined by anatomical connectivity among cells that have similar responses (Ko et al., 2011; Cossell et al., 2015; Lee et al., 2016). Viewing stimuli of a specific orientation during locomotion drives the orientation-specific subnetwork to increased firing because of disinhibition. Such an increase in synaptic activity would be expected to produce Hebbian plasticity specifically within that subnetwork, resulting in stronger connections among them that would continue to drive stronger responses even after the disinhibition was no longer present. Note that the changes in our experimental measurements using both intrinsic signal imaging and calcium imaging reflect the plasticity that was induced by the MLR-VIP-SST disinhibitory circuit, but they do not result from the activity of this circuit during the measurements, which were made in anesthetized animals in which the circuit is not active.

In neurons that showed an increase in Ca^2^+ responses to the experienced orientation with unchanged responses to other orientations, the measured preferred orientation consequently shifted toward the experienced orientation. This finding may reflect in part the fitting of tuning curves to coarsely sampled responses, at 45-degree steps in this experiment.

### Previous reports of enhancement

Our results from intrinsic signal recordings are generally in agreement with the previous reports in which recordings were made using VEPs (Frenkel et al., 2006; Cooke and Bear 2010), including observations of stimulus-specificity, requirement for NMDA receptor activation, persistence of enhancement after the sessions of visual stimulation were ended, and the degree of enhancement in mice older than P60. However, a critical difference between the present intrinsic signal and the earlier VEP studies is that in the latter animals were restrained during the daily exposure to visual stimuli, whereas we found no significant enhancement when the animals’ movements were restricted.

Several factors may contribute to this difference. First, different techniques were used to measure responsiveness. Intrinsic signal imaging used in this study is completely non-invasive and has been shown to produce stable read-out of visual cortical responsiveness over weeks (Kaneko and Stryker 2014). In contrast, chronic VEP recording requires electrode implantation right into the brain tissue at the center of interest, which could induce inflammatory responses including reactive astrocytes. Such disruptions can result in an increase in excitability of pyramidal cells resulting from dysfunction in astrocytes for microenvironment homeostasis and/or maintenance of GABAergic inhibition (reviewed by Robel and Sontheimer, 2016). This increased excitation-inhibition balance may induce abnormally high degree of plasticity in adult brain (reviewed by Bavelier et al., 2010).

Second, the VEP studies focused on layer 4 (L4), whereas the present study’s intrinsic optical signals reflect activity in L2/3 more strongly than in L4, with a significant contribution from L4 (Trachtenberg et al., 2000), and our chronic Ca^2^+ imaging was performed entirely on L2/3 cells. Perceptual learning has been shown to have different effects on L2/3 and L4 pyramidal neurons in mouse V1 (Makino and Komiyama 2015). L2/3 neurons acquired a new response pattern suggestive of anticipatory response while L4 neurons did not. This change in L2/3 was accompanied by increased excitatory drive from of top-down inputs. In rats, L2/3 excitatory cells in S1 showed tonic activation pattern that was longer in duration and higher in magnitude than L4 cells during active tactile discrimination, presumably also resulting from the influence by top-down inputs (Krupa et al., 2004). Similarly, after visual discrimination training in rhesus monkeys, sharpening of orientation tuning curve was observed in supragranular layer but not in L4 (Schoups et al., 2001).

In addition to laminar differences, intracortical microcircuits that participate in the enhancement of responses in L4 may be distinct from locomotion-induced response enhancement that we observed. A recent report has implicated the activity of parvalbumin-expressing (PV+) GABAergic cells in L4 enhancement (Kaplan et al., 2016). In contrast, as described above, a disinhibitory circuit comprised of VIP cells and SST cells plays a key role in control of the gain of visual responses and in facilitation of plasticity by locomotion in adult visual cortex, whereas PV neurons do not show consistent responses to locomotion (Fu et al., 2014, 2015).

### Outstanding questions about the role of locomotion and cortical plasticity

In the present experiment, the changes that were observed depend on locomotion. However, locomotion may be just one of many ways of activating the same circuit to enhance plasticity. While using locomotion to activate VIP cells in mouse V1 is particularly convenient for our experiments, recent findings indicate that “top-down” inputs from frontal cortex project to L1 of V1 to activate a similar disinhibitory circuit that almost certainly includes the same VIP cells during perceptual learning (Makino and Komiyama, 2015; Zhang et al., 2016). This idea is also consistent with many previous observations that responses in V1 were enhanced and visual behavior tasks were improved by pairing visual stimulation with manipulations that activate cholinergic inputs, such as application of cholinergic agonists and electrical or optogenetic activation of the basal forebrain (reviewed by Gu, 2003; Kang et al., 2014). Some recent studies have found that pupil dilation, often used as a measure of arousal, can be associated with changes in cortical responses similar to those produced by locomotion, but arousal is also activated by different systems with different effects on cortical activity (Reimer et al., 2014; Vinck et al., 2015).

We do not know how general is the stimulus-specific plasticity induced in V1 by locomotion, but preliminary reports do suggest an effect in humans (Lunghi and Sale, 2015; Bullock et al., 2016).

## ACKNOWLEDGEMENTS

This work was supported by NIH grants R01EY02874 and T32MH089920 and by the Simons Collaboration on the Global Brain project 325295. MPS is a recipient of the Research to Prevent Blindness Stein Innovator Award. We thank the members of the Stryker laboratory for assistance with and criticism of these experiments. The authors declare no competing financial interests.

